# coveR: An R package for processing Digital Cover Photography images to retrieve forest canopy attributes

**DOI:** 10.1101/2022.01.13.475850

**Authors:** Francesco Chianucci, Carlotta Ferrara, Nicola Puletti

## Abstract

1. Digital Cover Photography (DCP) is an increasingly popular tool for estimating canopy cover and leaf area index (LAI). However, existing solutions to process canopy images are predominantly tailored for fisheye photography, whereas open-access tools for DCP are lacking.
2. We developed an R package (*coveR*) to support the whole processing of DCP images in an automated, fast, and reproducible way. The package functions, which are designed for step-by-step single-image analysis, can be performed sequentially in a pipeline, while also allowing simple implementation for batch-processing bunches of images.
3. A case study is presented to demonstrate the reliability of canopy attributes derived from *coveR* in pure beech (*Fagus sylvatica* L.) stands with variable canopy density and structure. Estimates of gap fraction and effective LAI from DCP were validated against reference measurements obtained from terrestrial laser scanning.
4. By providing a simple, transparent, and flexible image processing procedure, *coveR* supported the use of DCP for routine measurements and monitoring of forest canopy attributes. This, combined with the implementability of DCP in many devices, including smartphones, micro-cameras, and remote trail cameras, can greatly expand the accessibility of the method also by non-experts.

## 1 INTRODUCTION

Accurate estimates of forest canopy attributes like canopy cover leaf area index (LAI) are central for a wide range of studies, including hydrology, carbon and nutrient cycling, and global change. As direct measurements are costly, time-consuming, and often destructive, indirect optical methods have been widely used in forest canopy research and monitoring, as documented by many published reviews (Breda, 2003; Chianucci, 2020; Jonckheere et al., 2004; Yan et al., 2019). Hemispherical photography has been amongst a pioneering method in forest ecology, with first applications dating back more than half a century ago (Anderson, 1964; Evans & Coombe, 1959), while its long-lasting use has been supported by the progress in digital photography and image processing technology (Chianucci, 2020).

Notwithstanding these improvements, some significant obstacles to the adoption of hemispherical canopy photography are:

1) the perceived sensitivity of the method to sky condition and camera exposure during image acquisition, which dramatically affects canopy gap retrieval (Macfarlane et al., 2000);

2) the complexity of fisheye image processing, which requires setting a circular mask, dividing the image into concentric annuli (zenith rings) and radial sectors (azimuth segments) to apply theoretical formulas to infer canopy attributes from the angular distribution of gap fraction (Chianucci, 2020).

All these factors hindered the standardization and comparability between image acquisition and procedures adopted using hemispherical photography, while also limiting the accessibility of the method by non-experts (Yan et al., 2019; Chianucci, 2020).

As an alternative to hemispherical photography, digital cover photography (DCP) has emerged in the last fifteen years as a robust and reliable method to estimate canopy structure in forest stands (Macfarlane, Hoffman, et al., 2007; Macfarlane, Grigg, et al., 2007; Ryu et al., 2011, 2012; Chianucci & Cutini, 2013; Lang et al., 2017; Chianucci et al., 2021). Compared with circular fisheye images, where only a fraction of the image pixels is used to sample the whole canopy, DCP uses all the image pixels to sample a narrow field of view (FOV), which brings several advantages. First, the high resolution (high number of pixels to sample a restricted portion of the canopy) and the uniform sky luminance yield few mixed pixels (Chianucci, 2016), which makes DCP relatively insensitive to sky condition (Macfarlane, Hoffman, et al., 2007), camera exposure (Macfarlane et al., 2014) and thresholding (Macfarlane, 2011). In addition, the use of a normal camera with restricted FOV increases the flexibility of DCP compared to other optical methods, which allowed to use this method for many purposes, including within-clumping correction (Macfarlane, Hoffman, et al., 2007; Macfarlane, Grigg, et al., 2007), isolated and urban tree measurements (Chianucci et al., 2015), phenological monitoring (Chianucci et al., 2021; Lang et al., 2017), post-disturbance canopy recovery assessment (Toda et al., 2018).

So far, the main limitation for a routine use of DCP is that most of the available open-access solutions to process canopy images have been limited to fisheye photography (see https://canopyphotography.wordpress.com/2021/12/06/open-access-tools-for-canopy-image-processing). Therefore, an effective step towards making DCP operational is to develop an open-source tool to reliably and efficiently process these images.

In this contribution, we introduced “*coveR*”, an R package to perform all the processing steps required to analyse and estimate canopy attributes from DCP images.

## 2 Introduction to Digital Cover Photography

The DCP method is based on digital images acquired below the canopy and oriented at the zenith, using a narrow lens (typically 30° FOV), to achieve a compromise between vertical looking and canopy sampling size. The restricted FOV yields high resolution close to the zenith, which allows separating the contribution of large, between-crowns gaps (*gL*) to total gaps (*gT*) in DCP images (Figure 1).

**Figure 1.**
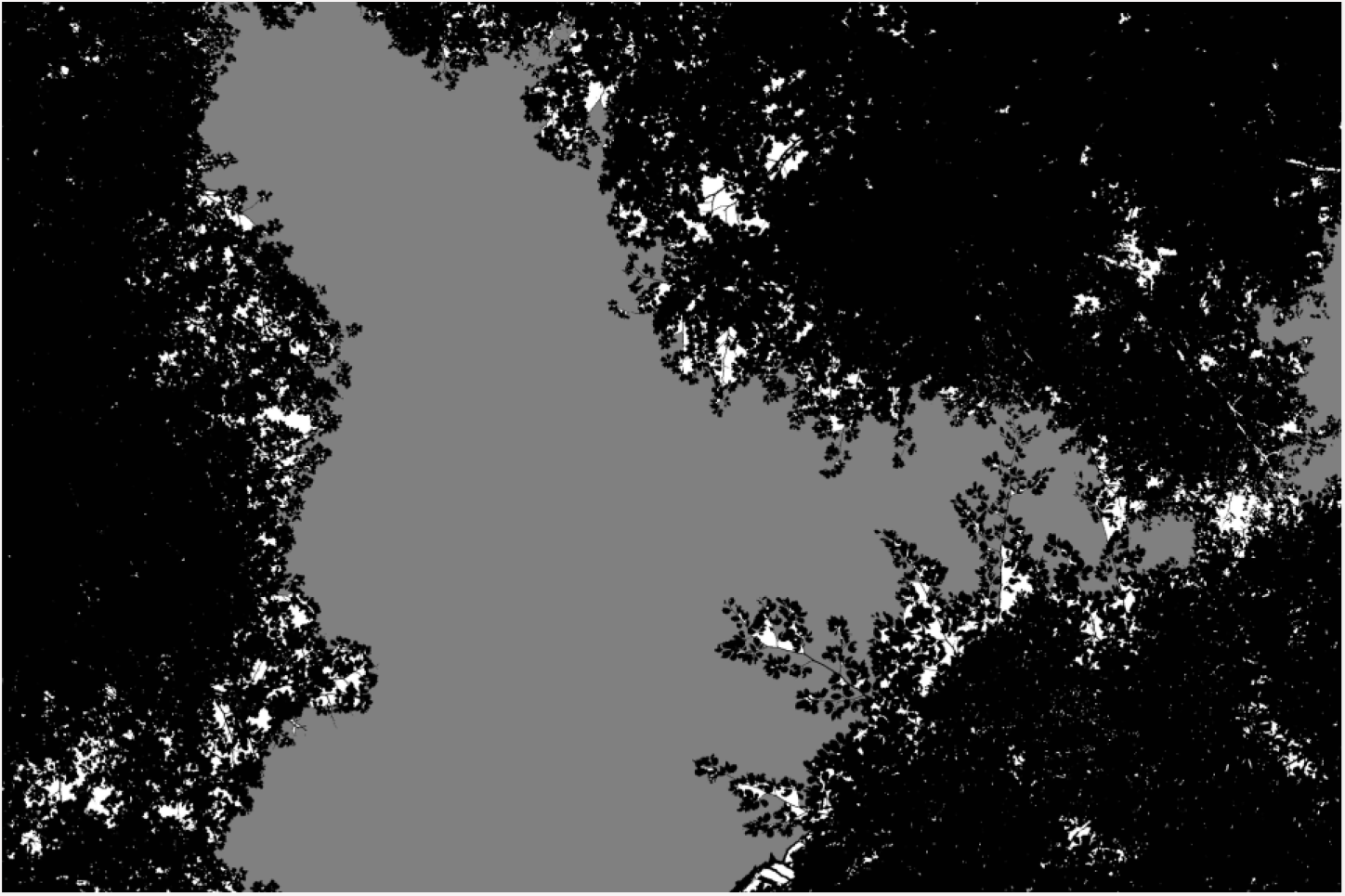
An example of a cover image that has been classified in canopy (black), large, between-crowns gaps (grey) and small, within-crown gaps (white). The image is obtained using the canopy_raster() function of the coveR package.

From these values, it is possible to estimate canopy cover attributes like foliage cover (*FC*), which is defined as the complement of gap fraction (*GF*), and crown cover (*CC*), which is defined as the fraction of the pixels that complement between-crowns gaps. Crown porosity (*CP*) is also calculated as the fraction of gaps within crown boundaries:

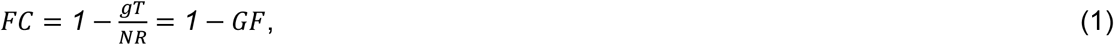

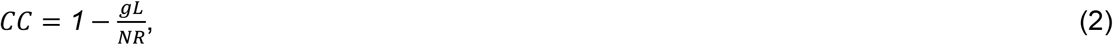

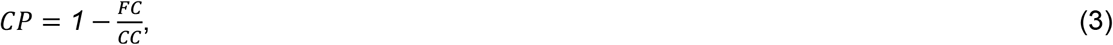

where *NR* is the total number of pixels in the image. From these canopy cover attributes, it is then possible to estimate effective (*Le*) and actual (*L*) LAI, by assuming an extinction coefficient at the zenith (*k*):

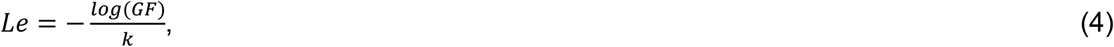

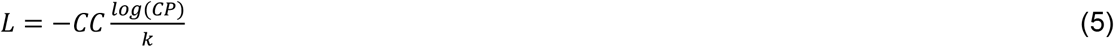

Finally, the clumping index (*CI*) is calculated as the ratio of *Le* to *L*:

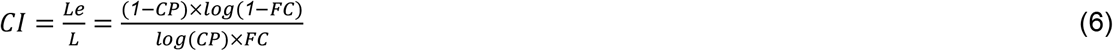

For details in the adopted equations, the reader is referred to Macfarlane, Hoffman, et al., (2007).

## 3 Overview of the coveR package

The key steps to process DCP images are basically:

i) import an image and select a single channel (typically the blue band);

ii) apply a method to classify the single channel pixels into canopy and sky;

iii) apply a method to further classify sky regions into large (between-crowns) gaps and small (within-crown) gaps;

iv) apply a modified version of the Beer-Lambert law (Macfarlane, Hoffman, et al., 2007) to estimate canopy attributes from classified images (see Equations 1-5 above).

Three additional considerations must be drawn when dealing with canopy image processing.

- Firstly, a strong advantage over other indirect methods lies in the permanent nature of digital images, which allows the inspection of the quality of acquired and processed data (Jonckheere et al., 2004). For this reason, an ideal tool for canopy image analysis would allow accessing the intermediate steps and output images generated from the processing workflow.
- With specific consideration of DCP images, a crucial point of the analysis is the classification of large gaps based on their size. While some manual approaches have been proposed for interactively labelling large gaps (Macfarlane, Hoffman, et al., 2007; Chianucci & Cutini, 2013), automated and objective methods are desirable for making the analysis reproducible and straightforward.
- The restricted FOV implies that a large number of DCP images need to be collected to achieve a statistically representative forest sampling (Chianucci, 2020). Therefore, an efficient tool for DCP would allow fast image processing time and simple batch-processing images capability.

To address the above considerations, *‘coveR’* uses the functionality of *‘raster’* package (Hijmans, 2021), which ensures faster processing of images not otherwise possible when dealing with other image formats in the R environment. In addition, the package is optimized for simple pipeline implementation using the ‘*%>%*’ functionality of ‘*magrittr*’ package (Bache & Wickham, 2020) and also for batch-processing images using the *‘map()’* functionality of the *‘purrr*’ package (Henry & Wickham, 2020), as illustrated in the Application section.

The package features the following key functions, which are ordered sequentially:

1. ‘*open_blue*’: imports an image and selects the channel for further analyses;
2. ‘*thd_blue*’: thresholds the selected image channel and returns a binary image;
3. ‘*label_gap*’: segments each sky region of the binary image and returns a labelled image;
4. *‘extract_gaps’*: calculates the size of each canopy and gap region;
5. *‘get_canopy’*: applies theoretical formulas to classify gaps based on their size, and retrieves canopy attributes from classified large and total gap fraction.
6. *‘canopy_raster’*: creates an image of the resulting classification of canopy, large and small gaps.

The processing workflow is graphically illustrated in Figure 2 and described in detail in the next sections. The key R package dependencies for image manipulations are: *raster* (Hijmans, 2021), *rasterDT* (O’Brien, 2020) and *EBImage* (Pau et al., 2010).

**Figure 2.**
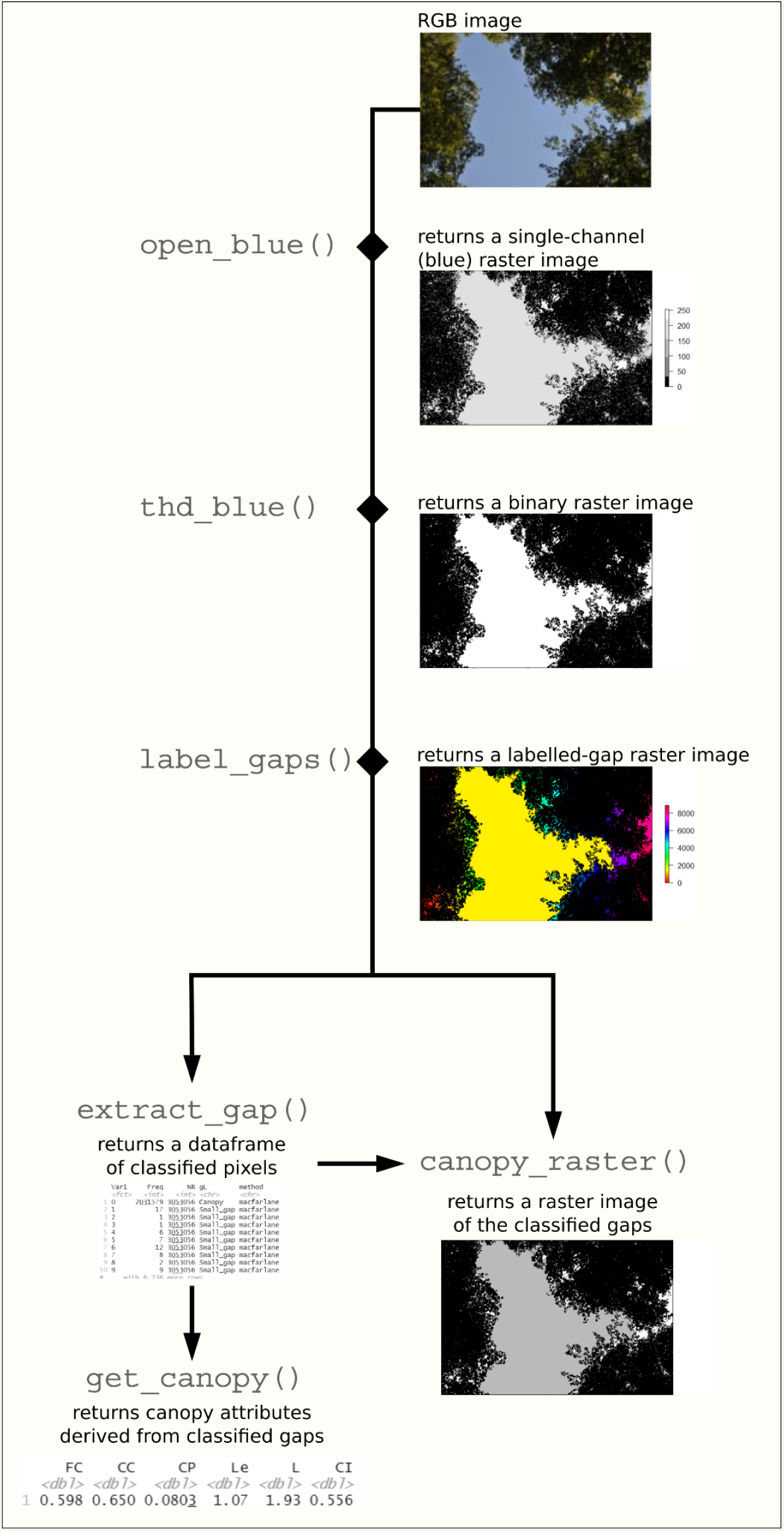
Illustrative example of the coveR package image processing workflow.

### 3.1 Import a cover image

The *open_blue()* function allows importing an image in the R environment as a raster file, using the *raster()* functionality of the *raster* package (Hijmans, 2021). The function requires specifying the number of the band corresponding to the blue channel (which by default is set as 3 in RGB images). The blue channel is generally preferred in canopy image analysis, as this portion of the electromagnetic spectrum allows best contrast between canopy and sky, particularly when images are acquired in diffuse sky conditions (Chianucci, 2020). This is well illustrated in Figure 3.

**Figure 3.**
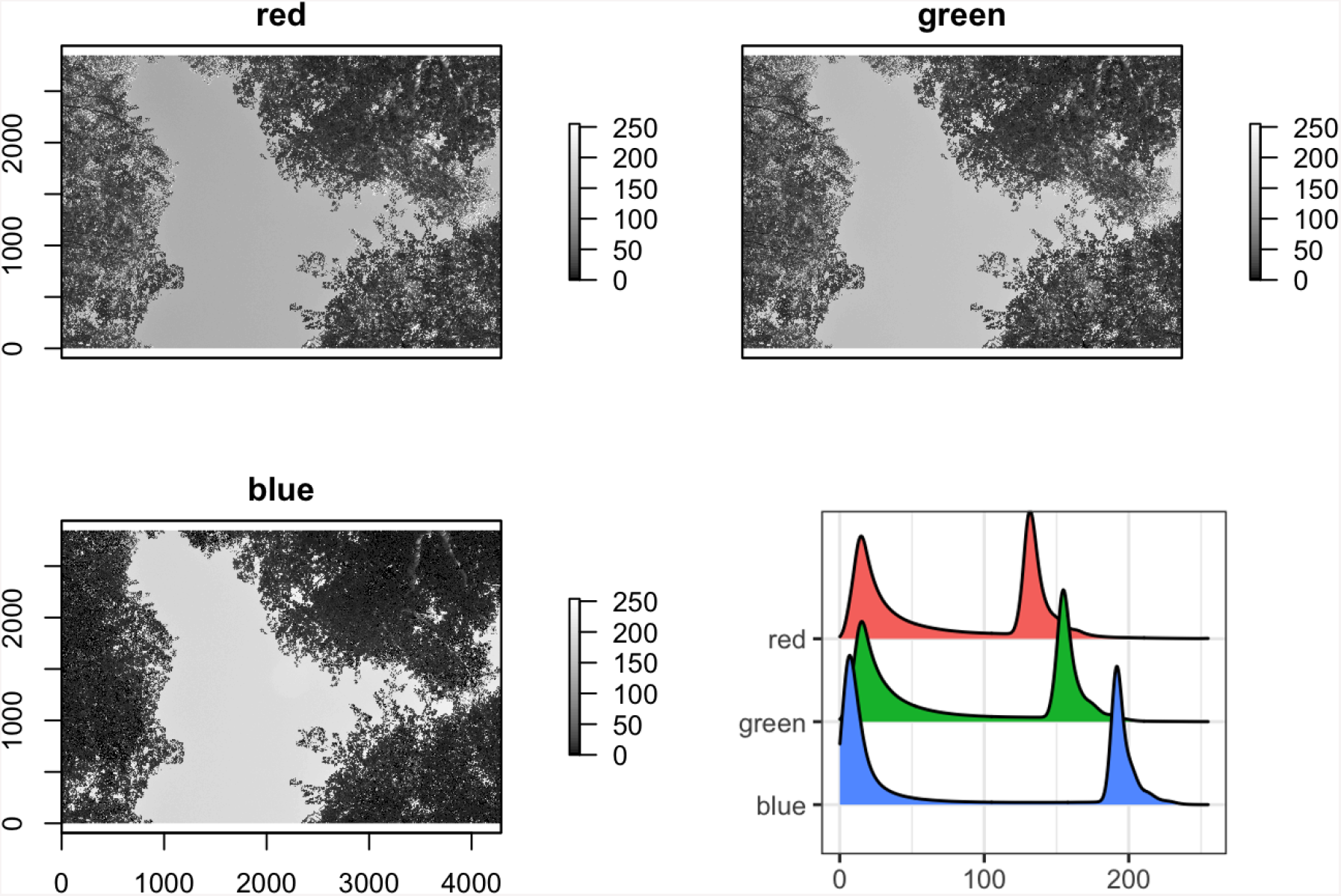
Greyscale version of the red, green and blue bands of a DCP image. In diffuse conditions, the sky is saturated in the blue band, and thus it appears brighter, while foliage has large light absorption, and thus appears darker, yielding a high contrast (high dynamic range) in the blue channel histogram.

### 3.2 Threshold an image

The *thd_blue()* function allows to classify the image channel considered and returns a binary raster image of canopy and gap pixels. It uses the *auto_thresh()* functionality of *autothresholdr* (Landini et al., 2017) to define a single thresholding method out of 17 available ones. The *autothresholdr* package is a wrapper of the ImageJ (Schneider et al., 2012) plugin ‘Auto Threshold’ (Landini et al., 2017). The default method was set to *‘Minimum*’, as it has been already proven effective in canopy images by Glatthorn & Beckschäfer (2014). Alternatively, the ‘*Otsu’* method has been often used in canopy image processing (e.g. Pueschel et al., 2012; Grotti et al., 2020).

### 3.3 Label gap regions in the binary image

The *label_gap()* function requires a binary raster image of canopy (0) and gap (1) pixels. The function then assigns a numeric label to each distinct gap region (where a region is a connected set of gap pixels). The output is a raster image, whose maximum value corresponds to the number of gap regions. The function uses the functionality of *bwimage()* of the *EBImage* package (Pau et al., 2010) to segment gap pixels.

### 3.4 Classify labelled gaps based on their size

The *extract_gaps()* function allows to classify each labelled gap region as large, between-crowns gap (‘Large_gap’) or small, within-crown gap (‘Small_gap’) based on their pixel size. The function allows users to select between two automated methods to classify large gaps. The default one (*method*=*‘macfarlane*’) classifies large gaps as those greater than a fraction of the image area; the default value is set to 1.3% (Macfarlane, Hoffman, et al., 2007; Macfarlane, Grigg, et al., 2007). Another method (*method=‘alivernini’*) defines a large gap threshold based on the statistical distribution of gap size of the image (Alivernini et al., 2018):

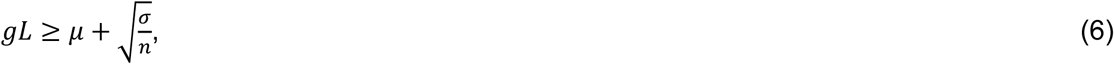

where µ and σ are the mean and standard deviation of image gaps (in pixels), *n* is the number of gap regions. Compared to the former, this method is density dependent, as the large gap threshold is self-calibrated on the actual canopy image analysed.

The output of the function is a dataframe with five columns, indicating:

– the image object (column *‘Var1’*). *Var1*=0 indicates canopy pixels, *Var1*>0 indicates distinct gap regions;
– the pixel size of each image object (column *‘Freq’*);
– the total image size (column *‘NR’*);
– the classification of each gap region (column *‘gL’*) in *‘Large_gaps’* or *‘Small_gaps’*. For *Var1*=0, *gL* =*‘Canopy’*;
– the method used to classify large gaps (column *‘method’*).

### 3.5 Retrieve canopy attributes from classified gaps

The *get_canopy()* function takes the dataframe of classified gaps as an input and returns a dataframe of the canopy attributes calculated from Equations 1-5 of the analysed image. The calculation of LAI requires setting a zenith extinction coefficient *k*, which by default is set to 0.5 (corresponding to a spherical leaf inclination angle distribution assumption; Pisek et al., 2013). The *k* parameter can be alternatively derived from measured leaf inclination angles (see Chianucci et al., 2018) or from existing datasets (Chianucci et al., 2018; Pisek & Adamson, 2020).

### 3.6 Export a classified cover image

The *canopy_raster()* function allows the user to return a raster of the classified image gaps (see an example in Figure 1), with assigned values 0, 0.5, and 1 to canopy, large gaps and small gaps pixels, respectively. The input arguments are the labelled image generated from *label_gaps()* and the dataframe of classified gaps generated from *extract_gap()*. The raster image derived from the function can also be exported, for example using the *writeJPEG()* function of *jpeg* package (Urbanek, 2021).

## 4. Application

### 4.1 Case study

To demonstrate the reliability of canopy attributes derived from *coveR*, we used a dataset of 315 cover images acquired in seven 0.25 ha pure beech (*Fagus sylvatica* L.) forest stands sampled in central Italy (Chianucci, 2020). The stands showed differences in canopy structure, resulting from different silvicultural regimes, with an effective *Le* estimated from TLS ranging between 1.3 and 5.3. Estimates of gap fraction (*GF*) and effective leaf area index (*Le*) from DCP were validated against reference measurements obtained with terrestrial laser scanning (TLS; Grotti et al., 2020). In each stand, nine TLS scans were acquired along a grid of sampling points, using a phase-shift FARO Focus 3D X130 laser scanner (Faro Technologies Inc., USA).

Given the different FOVs between the two methods, five DCP images per scan point were collected, with the first image acquired in the same TLS point location, and the other images acquired at four cardinal directions, within 5 m from each scan point. Reference canopy attributes were obtained from intensity images derived from raw TLS scan data, by setting a zenith angle range of 0-75° and 5 zenith bins, each 15° in size. Gap fraction was calculated for each zenith bin using (Otsu, 1979) multi-thresholding method, and *Le* was then derived by integrating the angular gap fraction using Miller’s (Miller, 1967) solution of the Beer-Lambert law (for details, see (Grotti et al., 2020). DCP images were collected using a Nikon D90 digital reflex equipped with a Nikkor 50 mm 1:1.8 D fixed lens and exported as 12MP in-built JPEGs. The blue channel of the DCP images was then selected and processed using the default functions’ arguments of the *coveR* package, with the exception of the extinction coefficient *k*, which was set to 0.85, based on a previous study (Chianucci, 2020).

Given the different FOVs, GF was firstly compared between the two methods considering only a zenith angle range of 0°-15° in TLS, and the nine aligned DCP-TLS sampling points. *Le* was then compared by averaging estimates from the five DCP images around each TLS scan point data. Results indicated that DCP images processed with *coveR* yielded accurate estimates of gap fraction, compared with intensity TLS images (Figure 4):

**Figure 4.**
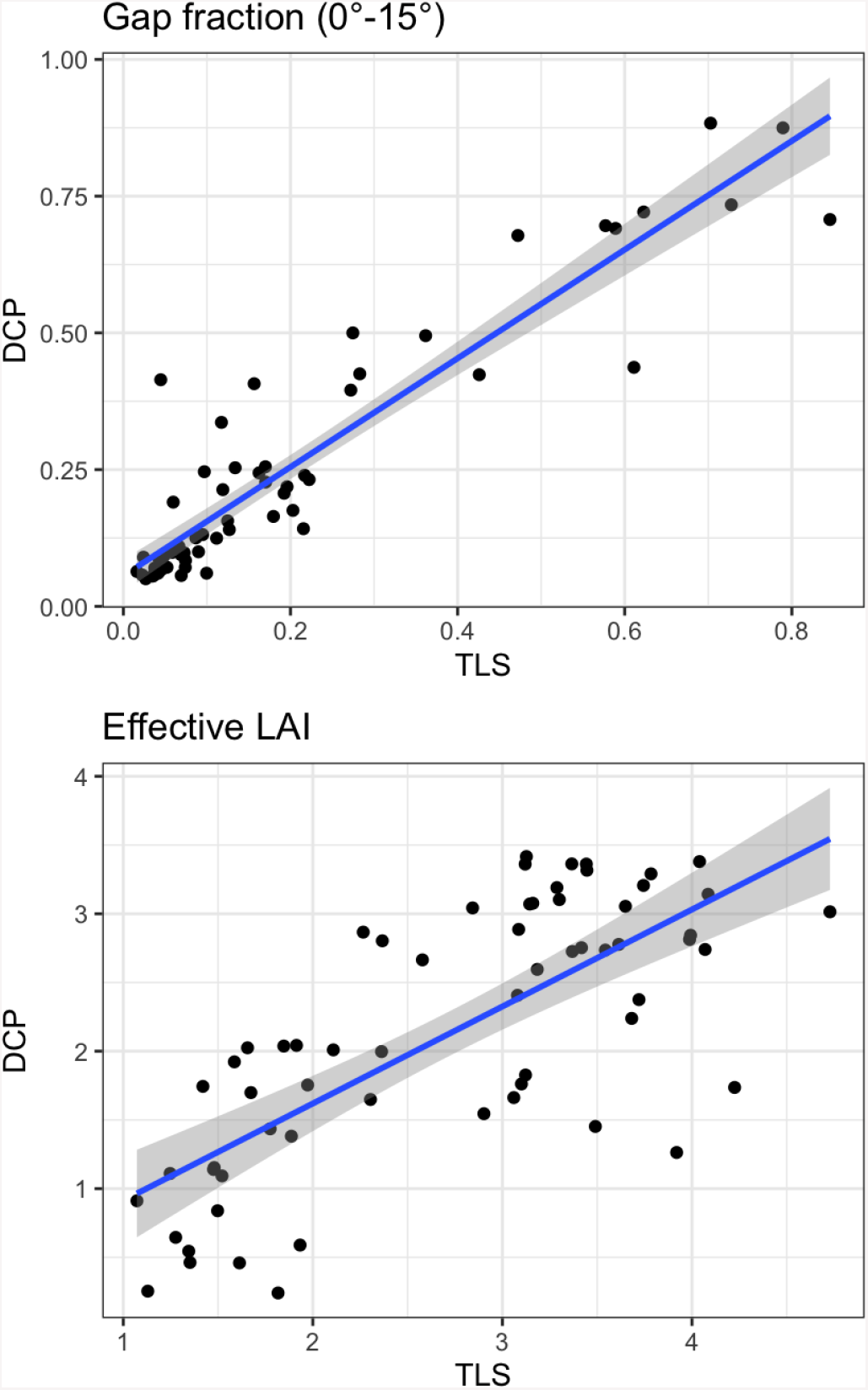
Left: Comparison between gap fraction estimates obtained from DCP and TLS at 0°-15° zenith angle range. Right: Comparison between effective leaf area index estimates obtained from DCP and TLS at their different FOVs.

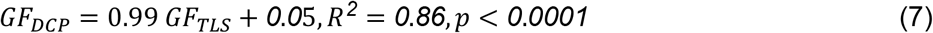

With regards to *Le*, the DCP method confirmed a good agreement with TLS, despite the differences in FOVs and theoretical formulas used between the two methods:

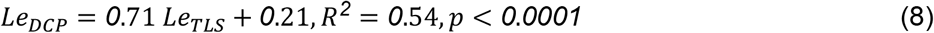

### 4.2 Optimizing coveR usage for single and batches images

The functions 1-5 implemented in the *coveR* package can be performed sequentially, using the pipe (*%>%*) operator (see the example below, considering the default arguments):

*open_blue() %>% thd_blue() %>% label_gaps() %>% extract_gap() %>% get_canopy()*

Given the different data outputs, the pipeline can be also splitted to separate raster from dataframe outputs. In such a case:

*open_blue() %>% thd_blue() %>% label_gaps()*

will generate raster outputs,

*extract_gap() %>% get_canopy()*

will generate dataframe outputs.

In addition, functions 1-5 can be also used for batch-processing bunches of images. In this case, we suggest setting the working directory (‘pathname’) where the images are stored, and using the *map()* functionality of *purrr* package (Henry & Wickham, 2020), as in the example below (considering JPEG images and the default function arguments):

*files<-dir(pathname, pattern=‘\**.*JPEG’)*

*files %>%*

*purrr::map(open_blue) %>%*

*purrr::map(thd_blue) %>%*

*purrr::map(label_gaps) %>%*

*purrr::map(extract_gap) %>%*

*purrr::map(get_canopy) %>%*

*dplyr::bind_rows()*

## 5 Conclusions

Digital cover photography has become an increasingly popular method to quantify canopy cover and leaf area index in forest stands. From an end-user’s perspective, the use of a normal camera with restricted view brings several advantages, as it allows implementing the DCP method to standard cameras, smartphones (Mora et al., 2016; De Bei et al., 2016), micro-cameras (Kim et al., 2019), and remote trail cameras (Chianucci et al., 2021), with significant reduction of equipment costs. This, combined with the availability of open-access digital image processing tools, can effectively extend the accessibility of the method to a wider audience. In this line, the *coveR* package provides a new, easy-to-use and out-of-the-box open solution for implementing DCP, making the process fast, robust, automated, and transparent.

From an operational point of view, the key features of the *coveR* package are:

– the embedding of *raster* functionality, which makes the single-image processing very fast (the full process requires about 5 seconds per image);
– the sequentiality of functions, which makes the package simple to use, while ensuring code conciseness and clarity, particularly when used in a pipeline (see examples in the paragraph Application);
– the ease implementation of the functions for batch-processing images (see examples in the paragraph Application).

While existing canopy photography protocols have mostly focused on fisheye photography, the *coveR* package can support the standardization of procedures for implementing DCP in long-term forest research and monitoring programs.

## Authors’ contributions

FC led the project, designed, and built the package, wrote original code and wrote the manuscript. CF supported manuscript draft revision and formatting, and package documenting. NP wrote the package functions, contributed to package building, supported manuscript draft revision.

## Data availability

The *‘coveR’* package can be installed from gitLab (https://gitlab.com/fchianucci/coveR). The available input DCP and TLS data used in examples are available from [repository link-after-article-acceptance].

